# ConsensusMetaDA: an R Package for consensus-based differential abundance analysis of metagenomic data

**DOI:** 10.1101/2025.11.03.686403

**Authors:** Manoharan Kumar, Matt A. Field

## Abstract

Differential abundance (DA) analysis is critical in metagenomics research for identifying microbial taxa associated with variable biological conditions. Despite numerous mature DA tools being available, they often generate disparate results, making it challenging to obtain consistent and biologically meaningful findings. Tools differ substantially in their input requirements, underlying statistical models, and filtering strategies, yet researchers typically rely on a single tool due to the practical challenges of integrating multiple algorithms with variable workflows and non-standardized output formats.

To address this challenge, we developed ConsensusMetaDA, an easy-to-use R package that integrates five widely-used DA tools (ALDEx2, DESeq2, EdgeR, metagenomeSeq, and ADAPT) into a streamlined three-function workflow. The package generates unified output reports and comprehensive visualizations including rarefaction curves, alpha/beta diversity analyses, taxonomic abundance plots, and Venn/UpSet diagrams to visualize algorithmic consensus and disagreement.

We benchmarked ConsensusMetaDA on well-characterized public datasets and simulated data, demonstrating that individual tools exhibit complementary strengths and weaknesses. On datasets with known null results, EdgeR and DESeq2 reported false positive differentially abundant OTUs while ALDEx2, metagenomeSeq, and ADAPT correctly identified none. Conversely, simulations revealed that ALDEx2, metagenomeSeq, and ADAPT frequently missed true positives that EdgeR and DESeq2 detected. Our consensus approach improves both sensitivity and specificity compared to any single tool and enables researchers to minimize either false positive or false negative rates based on study-specific priorities.

ConsensusMetaDA is freely available with comprehensive documentation at: https://github.com/FieldLabFNQOmics/ConsensusMetaDA

## Background

Metagenomic sequencing is widely used for environmental and human health studies. In human samples, empirical studies on microbiome data have shown that gut microbial diversity plays a significant role in food digestion, nutrient intake, and overall human health (1-4) while environmental samples uncover how microbial communities and their functional potentials vary across different conditions such as ecosystems, pollution gradients, climate zones, or treatment groups. For both applications, an increasingly common use of metagenomic is the identification of differentially abundant (DA) species associated with variable health or environmental conditions. There is a wide range of tools available for DA analysis, some of which were originally designed for RNA-Seq gene count analyses including Limma (5), DESeq2 (6) and EdgeR (7). More recently metagenomic-specific DA frameworks and algorithms have been developed including MetaDEGalaxy (8), metagenomeSeq (9), MaAsLin2 (10), ANCOMB-II (11), and ALDEx2 (12).

While the increasing number of tools is encouraging, recent benchmarking studies demonstrate a lack of concordance between leading algorithms (13). Further, running each tool can be challenging requiring specific formats for input file(s), usually a count table, sample metadata, and taxa details. Additionally, there are often numerous algorithm-specific steps involved in converting sequencing reads to the required input data as well as differences in the underlying models of the DA algorithms. For example, popular tools such as DESeq2 and edgeR are not specifically designed for metagenomic data in contrast to a tool like ALDEx2 which is designed to consider compositional variation in the data. These underlying differences drive the discordant results and suggest an optimal solution may be a consensus-based approach, with recent studies showing improvements in sensitivity and specificity across a variety of common sequencing applications including RNA-Seq and variant detection (14-16).

Despite the need, there are currently limited consensus-based options for metagenomic DA.

To address this issue, we have developed ConsensusMetaDA, an R package that incorporates five commonly used tools including DESeq2 (17), edgeR (7), metagenomeSeq (18), ADAPT and ALDEx2 (12). In addition to producing consensus-based DA results, ConsensusMetaDA generates common standard metagenome plots as well as Venn diagrams and UpSet plots to visualise the algorithmic overlap. ConsensusMetaDA is designed with simplicity in mind, running all analyses with three simple commands, ensuring robust and reproducible output less subject to the limitations of any single algorithm.

## Methods

Unless stated otherwise all data analyses were carried out in R version 4.4.2 (20) using Bioconductor. ConsensusMetaDA requires R packages including phyloseq (v1.46.0), ggplot2 (v3.5.2), edgeR (v4.4.2), DESeq2 (v1.46.1), ADAPT (v1.0.0), metagenomeSeq (v1.48.1) and ALDEx2 (v1.38.0). For all tools we employ default normalization methods. ConsensusMetaDA requires three files as input: a BIOM file, a metadata sample table consisting of ‘Sample’ and ‘Group’ columns, and a taxonomy table (**Figure 1A**). The workflow consists of three steps corresponding to OTU table construction (build_OTU_counts), differential abundance analysis (OTUs_multi_DA) and plot generation (OTU_plots) (**Figure 1B**).

**Figure 1:**
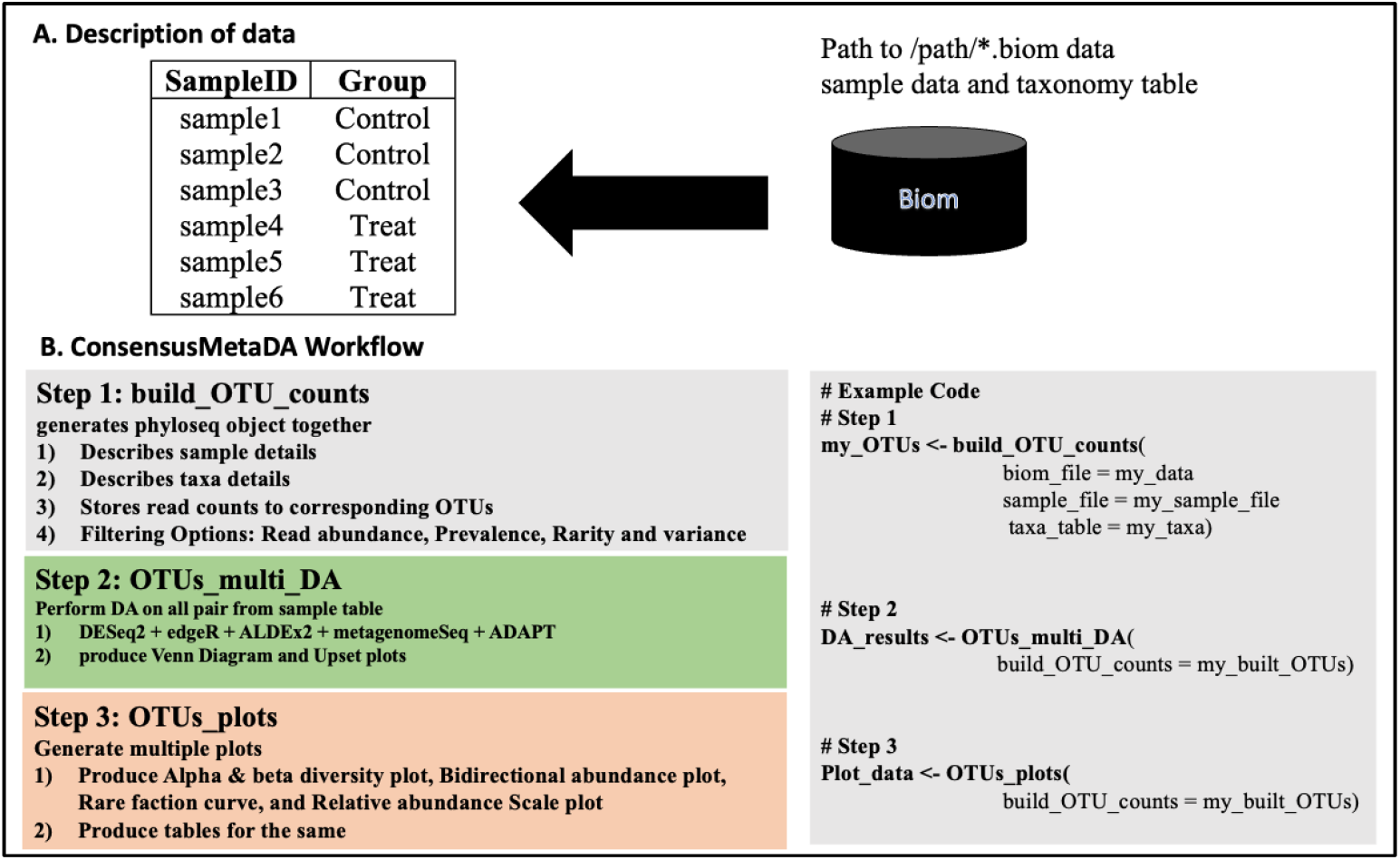
ConsensusMetaDA data requirements and workflow. A) ConsensusMetaDA requires three inputs including a BIOM file, a sample metadata table and taxonomy table. B) ConsensusMetaDA consists of three main functions: i) build_OTU_counts() converts user-supplied BIOM file and metadata file into a phyloseq object ii) OTU_multi_DA() runs all five algorithms and overlaps results iii) OTU_plots generates five plots (alpha and beta diversity, rarefaction curves, bidirectional DA OTU plot and proportional taxonomic bar graph) to visualize the data and results.

The details of the functions are:

1. **build_OTU_counts:** Generates summarized experiment from input BIOM file, metadata sample table and taxa table information. Currently taxa tables support seven taxonomic levels from Kingdom to Species. This function creates phyloseq object using the phyloseq package (v1.46.0) (21). The build_OTU_counts() function contains options for filtering OTU counts such as abundance_threshold, prevalence_threshold, rarity_threshold and variance_threshold, all of which can be overwritten as needed.
2. **OTUs_multi_DA:** This function performs DA analysis on all possible pairings of groups contained in the sample table for all five methods. For each comparison, it generates results for both individual algorithms and for the overlap of the algorithms with overlap plots (Venn/UpSet) allow users to visualise the level of concordance between the algorithms. The combined summary table including columns such as “mean_LogFC, “mean_Log-SD”, “edger_adj_p”, “deseq2_adj_p”, “ALDEx2_adj_p”, “metagenome_adj_p”, “ADAPT_adj_p”, “rank_sum”, “p_intersect” and “p_union” with the results sorted by rank-sum (22) to prioritise differentially abundant species shared amongst the algorithms. Supplementary **Table S1** describes the output columns and default parameters in detail.
3. **OTUs_plots:** The function generates five common metagenomics plots as PDF files: 1) Alpha diversity (Shannon index), 2) Beta diversity - Principal Coordinate Analyses (PCoA), 3) Rarefaction curve 4) Scaled abundance plots at various taxa levels and 5) Bidirectional abundance plot containing the most differentially abundant OTUs. These plots are all generated using the ggplot2 package (23).

Benchmarking was performed on both real and synthetic data to estimate false positive and false negative rates respectively. False positive rates were determined following a similar approach described in previous benchmarking studies (13) where two real world datasets (24, 25) were artificially split into separate groups representing a scenario where no statistically significant differences would be expected. Using this approach, each dataset was run ten times. False negative rates were determined using synthetic data generated with the SparseDOSSA (v0.99.6) (26), an R package which simulates metagenome data using a linear model to generate distinct groups. The package contains parameters to modify the scale of the differences including number of OTUs, proportion of DA species, sequencing depth and fold-change. We set fold-change to 10X, sequencing depth at 20,000 reads/OTU and total OTUs at 500, with all other parameters kept as default. We ran this simulation 100 times with randomly generated seed values and varied the proportion of DA OTUs from 5 to 100%.

## Results

To demonstrate the functionality of our tool, we processed a publicly available sediment dataset (https://figshare.com/articles/dataset/16S_rRNA_Microbiome_Datasets/14531724) (13, 27) with ConsensusMetaDA v1.0 using default parameters. build_OTU_counts() generated a matrix of 3,000 OTUs spread across 40 samples and two conditions. OTUs_multi_DA() ran the five DA algorithms, generating individual results, a combined summary table and a Venn diagram and UpSet plot to visualise algorithmic overlap (**Figure 2**).

**Figure 2:**
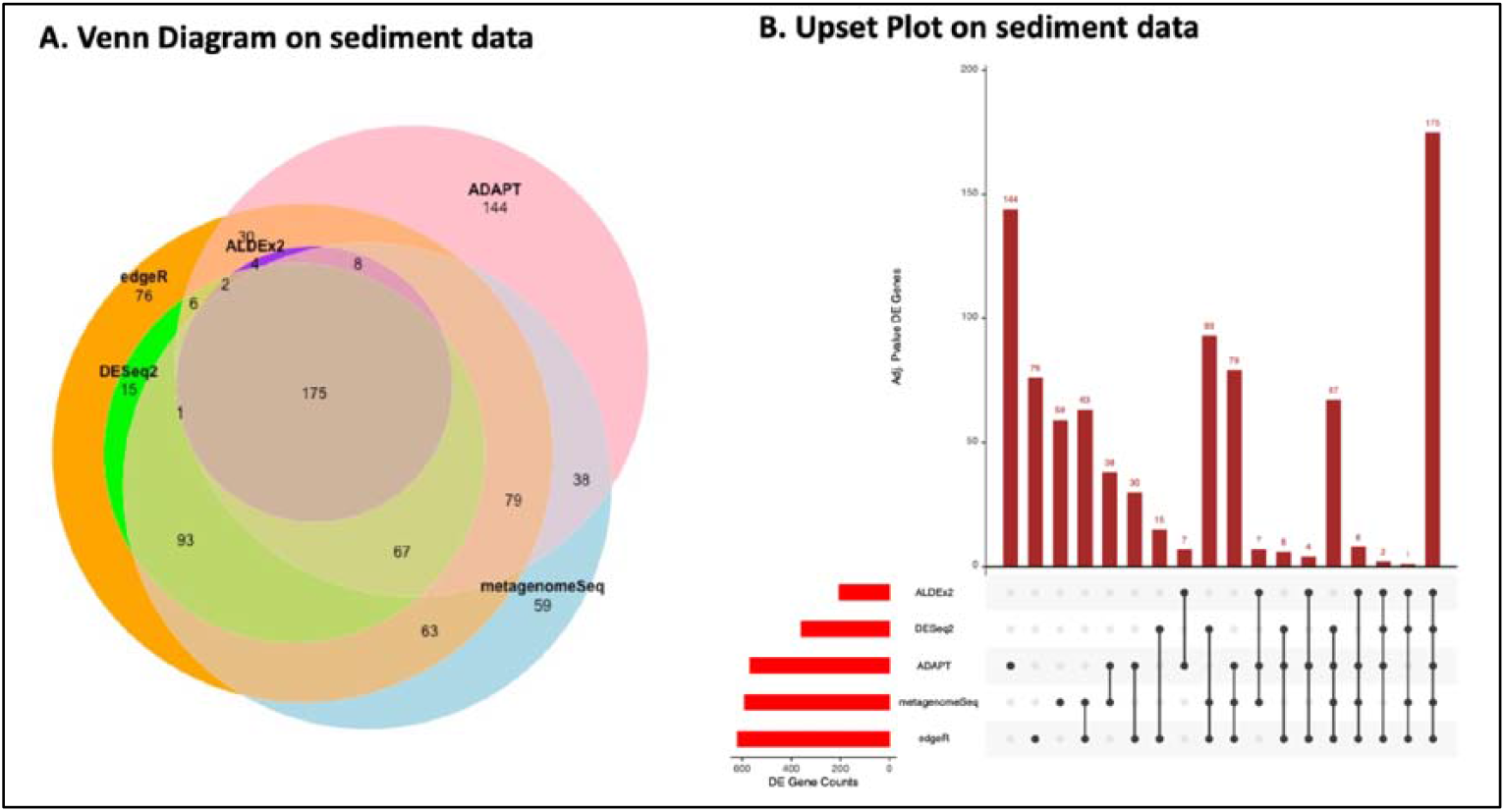
ConsensusMetaDA algorithm overlap using publicly available sediment data. By default, any OTU with an adjusted p-value of <0.05 was classified as DA OTUs and taken forward. The Venn diagram (A) and the Upset plot (B) visualise the concordance of all algorithms while additionally highlighting the large numbers of algorithm specific DA OTUs.

Overall, a total of 874 DA OTUs were detected, with only 175 detected by all five tools (20% of total) and 279 unique to a single tool (32% of total with a range of 0-144 per algorithm). Both DESeq2 and ALDEx2 did not detect any algorithm specific DA OTUs, while metageomeSeq, edgeR and ADAPT reported 59, 76 and 144 algorithm-specific DA OTUs respectively. A further 173 DA OTUs were reported by exactly two algorithms (20% of total), the largest number common to edgeR-metagenomeSeq (63 OTUs) followed by ADAPT-metagenomeSeq (38 OTUs). There were slightly more DA OTUs common to three algorithms (189 or ∼22% of total), the majority of which we detected by DESeq2-edgeR-metagenomeSeq (93 OTUs) and ADAPT-edgeR-metagenomeSeq (79 OTUs). Finally, a further 78 DA OTUs were detected uniquely by four algorithms (78 or 9% of total), the majority of which were not detected by ADAPT, generally found to be the most conservative algorithm. Collectively, these results demonstrate significant discordance across the five algorithms, justifying the utility of a consensus approach. Overall, the discordant levels observed are similar to previous benchmarking studies and not unexpected given the differences in each algorithm’s distribution model and hypothesis tested (**Table S2**).

To demonstrate the plotting capabilities, we subsetted ten samples from an additional publicly available 16S BIOM data consisting of samples spanning three conditions (ftp://ftp.microbio.me/emp/). We called the OTUs_plots function to generate all main plots for alpha and beta diversity, rarefaction curves, bidirectional plot of top DA OTUs and relative abundance plots (**Figure 3**). For diversity, the beta diversity plot demonstrates sample clustering by condition (**Figure 3A**) while the alpha diversity shows significant differences amongst the groups (**Figure 3B**). The taxonomic relative abundance plots show the relative abundance of the samples at the taxonomic level of Class (**Figure 3C**) while the bidirectional DA OTUs plots (**Figure 3D**) identifies the most significant DA OTUs. Finally, the rarefaction plots (**Figure 3E**) demonstrate variable levels of saturation with increasing data. By default, relative bidirectional DA OTUs plots and abundance plots are generated at all taxonomic levels.

**Figure 3:**
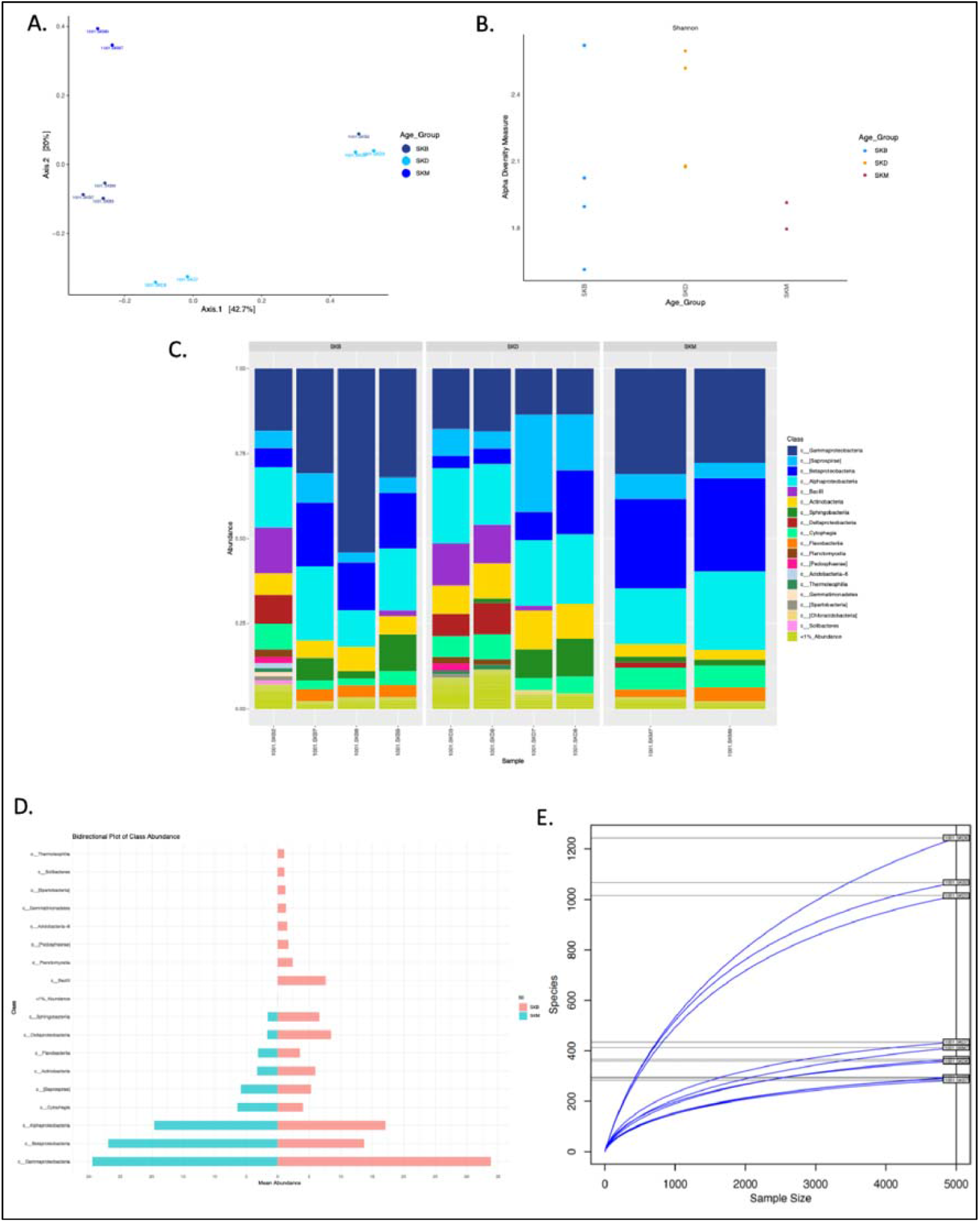
Five plots generated with the OTUs_plots. A) Beta diversity (Principal Coordinates Analysis) captures inter-sample diversity; B) Alpha diversity captures intra-sample diversity; C) Relative abundance taxonomic plot represents microbial distribution within samples and larger groups; D) Bidirectional plot of most significant DA OTUs shows mean abundance for each group shown; and E) Rarefaction curve shows saturation of microbial species across increasing sequences per sample.

To estimate false positive (FP) rates, we employed a similar approach from previous benchmarking studies (13) that artificially split a single data set into two groups where ideally no differences are expected. We performed this analysis across two publicly available data sets, namely COLD_HOT and Office (24, 25). Each data set was split ten times into two even groups and the number of DA OTUs calculated. Overall, only a small number of FP DA OTUs were reported for the five algorithms, with only edgeR and DESeq2 generating any FP DA OTUs (**Figure 4A**). Overall, DESeq2 had only a 1% FP rate for the COLD_HOT data set, while edgeR reported ∼5% FP rates for both datasets, indicating this is a larger problem for the algorithm.

**Figure 4:**
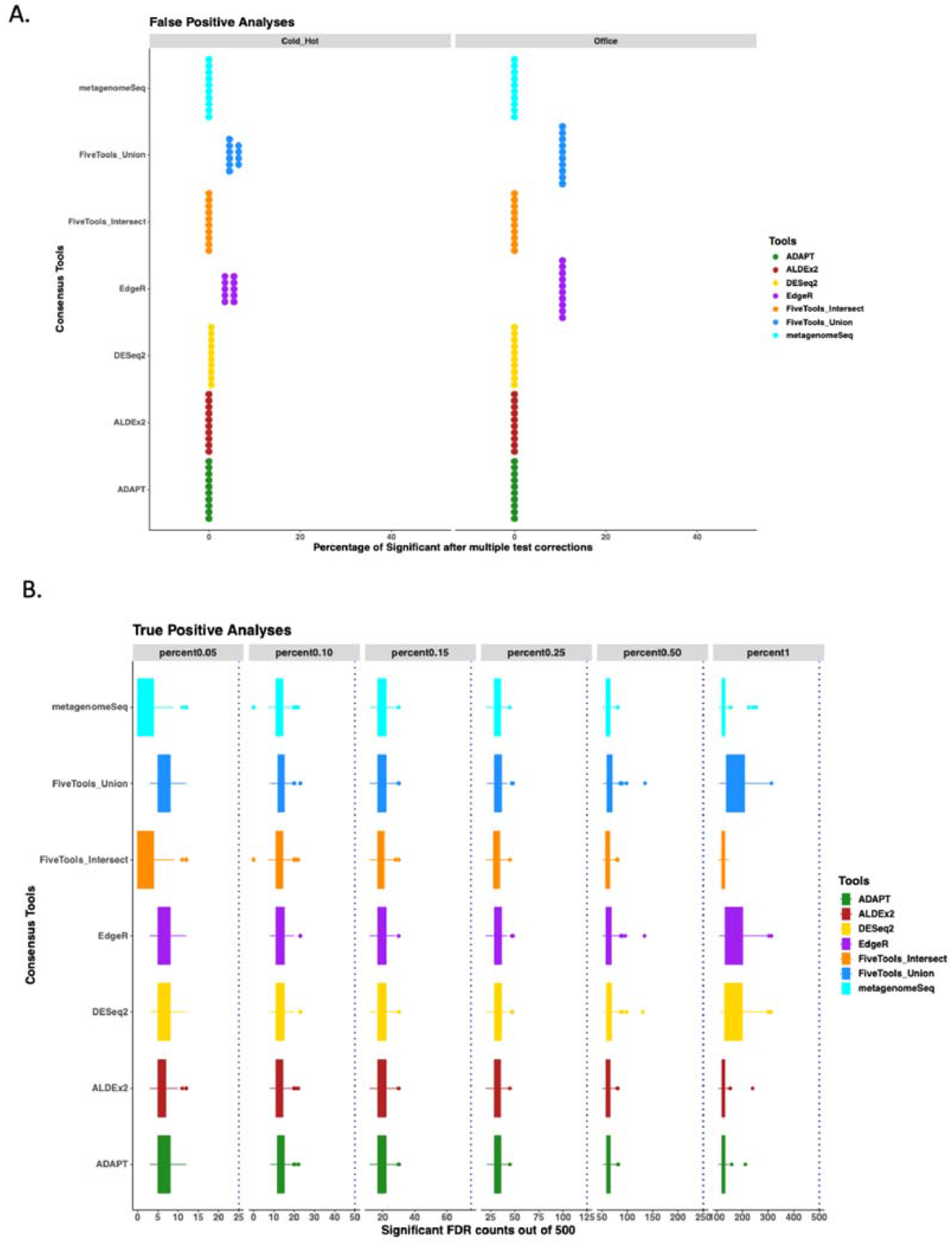
False positive and true positive analyses for the five individual algorithms and their union and the intersection. A) False positive rates were measured by artificially dividing a uniform data set into two distinct groups and running ten iterations. B) True positive rates were measures using synthetic data from SparseDOSSA R package varying the percentage of DA OTUs at 5%, 10%, 15%, 25%, 50% and 100% resulting in an expected 25, 50, 75, 125, 250 and 500 DA OTUs (represented by dotted line). We ran 100 iterations for each algorithm (plus the union and intersection of the five algorithms) with box and whisker plots representing the distribution.

To estimate true positive rates, we employed simulated data generated using the linear model from the SparseDOSSA R package. For this analysis, we ran 100 iterations across 500 OTUs with a set differential level of 10X and varying percentages of DA OTUs (5%, 10%, 15%, 25%, 50% and 100% resulting in an expected 25, 50, 75, 125, 250 and 500 DA OTUs) **(Figure 4B)**.

Overall, all algorithms miss significant numbers of DA OTUs, with average true positive (TP) rates ranging from 24-37% across all algorithms and permutations. There are also significant differences within the 100 iterations for each algorithm/permutation combination; for example, metagenomeSeq finds between 0-12 DA OTUs of the expected 25 at 5%, while DESeq2 find between 8-23 DA OTUs of the expected 50 at 10%. The performance differences become more pronounced where the proportion of real DA OTUs is increased. For example, with 100% of OTUs set as DA, edgeR and DESeq2 find an average of ∼50 additional DA OTUs relative to the other algorithms. Furthermore, the best TP rates are achieved taking the union of all algorithms, with five additional DA OTUs identified in the union relative to any single algorithm. Thus, if minimising false negatives is the project priority, the value of the consensus approach is clear.

## Discussion

Identifying differentially abundant biologically relevant OTUs is one of the most common applications in metagenomic studies (28-31). Despite the importance of this application, large-scale metagenomic benchmarking studies routinely demonstrate significant differences in the performance of algorithms with one study concluding a consensus approach is the optimal solution (13). The consensus approach is increasingly becoming standard practice within many common sequence-based analyses (14, 32) however it is not standard within DA OTU analysis due to a variety of factors. For example, each individual DA tool requires custom input files and further requires variable number of function calls to perform a standard metagenomic DA analysis producing the output tables and plots. For example, the ADAPT package requires a phyloseq object as input whereas the other tools accept count matrices. Some packages like DESeq2 run with a few simple commands whereas others like metagenomeSeq have over 80 functions available representing a significant learning curve. To remove such learning curves, ConsensusMetaDA was designed with simplicity in mind focusing on removing the specific R-packages requirements and replacing them with three simple intuitive commands for building the counts table, running the DA analysis and generating the relevant plots. Further, the entire workflow requires only three simple input files: a BIOM file, a sample/group metadata file and a taxonomy table. While customisation via additional parameters is possible, at its core ConsensusMetaDA is extremely simple and intuitive to use.

Individual DA algorithms generate semi-standardised metagenomic plots however there is little consistency across the algorithms in number and style of function calls and the number of customisable features and methods for generating individual plots. To address this, we generate the major standard metagenomic plots with one simple function call creating plots including alpha and beta diversity and rarefaction curves. Further, by default we generate plots for sample-level taxonomical abundance distribution and a bidirectional plot of the top DA OTUs for each taxonomic level, thus removing the need to run the plotting functions multiple times across the taxonomic levels of interest. Lastly we generate Venn diagrams and UpSet plots to visualise the overlap between the five algorithms allowing the users to decide on the set of OTUs to take forward based on the project priorities. Collectively the plots generated provide the user with an understanding of the overall structure and composition of the data.

While our results for FP and TP measurements highlighted small differences amongst the algorithms, a more typical result is represented by the analysis on the sediment data highlighted in Figure 2. In this analysis, the intersection of the five algorithms identified only 20% of the total DA OTU calls (175/874). At the other extreme, 32% of total DA OTUs (279/874) were called by a single algorithm, with the remaining roughly 48% reported by 2-4 algorithms. At the two extremes, the results from the intersection of the tools will be significantly enriched for true positives, while the single algorithm calls are enriched for false positives; however how to approach the 48% of the calls reported by 2-4 algorithms is unclear. Deciding whether to include these intermediate level DA OTUs is ultimately project dependent, with considerations around whether false positives or false negatives are more important to the underlying project goals. We recognise this and thus designed ConsensusMetaDA flexibility, by default offering users the ability to consider either the union or the intersection of the five algorithms. consider the results from either one algorithm or the union / intersection of the results. We further generate a single summary file containing all DA OTUs and a list of algorithms supporting the call allowing custom rule development. For example, custom rules could be DA OTUs found by three or more algorithms or those found by a specific algorithm and at least one more algorithm, etc. Ultimately the balance between precision and recall lies with the user.

We tested ConsensusMetaDA across datasets of variable size and composition to ensure reasonable run time and memory usage. While largely limited by the requirements of the underlying algorithms we have run datasets containing 5,000 OTUs and 500 samples in less than one hour using <32GB memory. This robustness is particularly important for metagenomic studies since there is a lot of variability in sample size, sequencing depth, and biological conditions. While sequencing technologies continue to increase read length and increase throughput, our tool’s run time is unlikely to change from this factor given the input requirement of a BIOM count matrix. Significant increases in number of samples and OTUs will increase run times however, with future versions potentially offering the ability to omit particularly time-consuming algorithms.

The current release of ConsensusMetaDA has several limitations that warrant attention. Currently, it relies on five specific tools (DESeq2, edgeR, ALDEx2, metagenomeSeq, and ADAPT) which may exclude taxa that could be identified by other improved tools as they are developed. Future releases of ConsensusMetaDA could incorporate additional tools to capture any improvements in this area with the flexible design lending itself to the integration of any number of additional tools. Another consideration is the expectation that the input BIOM is preprocessed having gone through QC and normalisation beforehand. In future releases, we aim to incorporate an optional automated preprocessing pipeline to enhance usability and make the entire workflow possible within our package. Finally, this package is primarily designed for 16S metagenomic sequencing data; further extending the tool to other types of microbial studies such as shotgun-based metagenomic data and functional abundance analyses will extend its’ utility. Despite these limitations, ConsensusMetaDA represents a meaningful contribution to the field, offering users a simple way to run disparate and complex individual DA OTU algorithms in a simple, unified framework.

## Conclusion

We created a new R package ConsensusMetaDA to perform consensus-based differential abundance analysis for metagenome data. Our results show consensus-based results provide the user with the ability to minimise either false positive or false negative rates depending on the project specific goals.

## Supplementary Details

Table 1: ConsensusMetaDA output column names and descriptions.

Table 2: DA Software included in ConsensusMetaDA.

## Acknowledgments

The authors are thankful to the people who tested all versions of this R package and the National Computational Infrastructure (Australia) for continued access to significant computation resources and technical expertise.

## Data Availability

The code is available at GitHub: https://github.com/FieldLabFNQOmics/ConsensusMetaDA

## Supplemental Data

**Table S1:** ConsensusMetaDA output column names and descriptions.

| ID | Description |
| --- | --- |
| LogFC | Mean Log Fold change for DESeq2 and edgeR |
| LogFC_sd | Mean Log Fold change for DESeq2 and edgeR standard deviation |
| padj_edgeR | edgeR adjacent p-value |
| padj_DESeq | DESeq adjacent p-value. |
| padj_ALDEx2 | ALDEx2 adjacent p-value. |
| padj_metagenomeSeq | MetagenomeSeq adjacent p-value |
| padj_ADAPT | ADAPT adjacent p-value |
| rank_sum | All three tools combined rank sum |
| p_intersect | Empirical Benjamini-Hochberg (BH) adjusted p-value for the between group. |
| p_union | Empirical Benjamini-Hochberg (BH) adjusted p-value for the between group. |

**Table S2:** DA Software included in ConsensusMetaDA.

| Tools | Input | Distribution | Language | Hypothesis test |
| --- | --- | --- | --- | --- |
| ALDEx2 | Counts | Dirichlet-multinomial | R-Bioconductor | Wilcoxon rank-sum |
| DESeq2 | Counts | Negative binominal | R-Bioconductor | wald |
| edgeR | Counts | Negative binominal | R-Bioconductor | Exact |
| MetagenomesSeq | Counts | Zero-Inflated Gaussian | R-Bioconductor | t-test/z-test |
| ADAPT | phyloseq | Left-Censored Normal Distribution | R-Bioconductor | Tobit Model |

